# Effects of rhein on bile acid homeostasis in rats

**DOI:** 10.1101/2020.01.29.925297

**Authors:** Zhong Xian, Jingzhuo Tian, Lianmei Wang, Yushi Zhang, Jiayin Han, Nuo Deng, Suyan Liu, Yong Zhao, Chunying Li, Yan Yi, Dunfang Wang, Jing Meng, Chen Pan, Aihua Liang

## Abstract

Rhein, the active ingredient of rhubarb, a medicinal and edible plant, is widely used in clinical practice. In this work, we investigated the alterations of 14 bile acids and hepatic transporters after rats were administered rhein for 5 consecutive weeks. There was no obvious injury to the liver and kidney, and there were no significant changes in biochemical indicators. However, 1,000 mg/kg rhein increased the liver total bile acid (TBA) levels, especially taurine-conjugated bile acids (t-CBAs), inhibited the expression of Farnesoid X receptor (FXR) and (bile salt export pump) BSEP mRNA, and upregulated the expression of (cholesterol 7α-hydroxylase) CYP7A1 mRNA. Rhein close to the clinical dose reduced the amounts of TBAs, especially unconjugated bile acids (UCBAs), and elevated the expression of FXR and multidrug resistance-associated protein 3 (Mrp3) mRNA. These results denote that rhein is not toxic and is safe to use at a reasonable dose and timing.

## Introduction

Bile acids play an important role in regulating the metabolism balance of lipids *in vivo* (1). They are converted from cholesterol in liver by a series of enzymes, and they maintain a dynamic balance through the uptake and efflux of hepatocellular transporter as well as the enterohepatic circulation (2). Some liver diseases and drug-induced liver injury are often accompanied by obstacles in the synthesis, metabolism and excretion of bile acids, potentially leading to the accumulation of bile acids. However, excessive amounts of bile acid results in hepatocyte injury, even liver cirrhosis and necrosis (3). The homeostasis of bile acid levels is closely related to the occurrence and development of liver diseases. Despite the limitations of clinical chemistry in sensitivity and specificity at the early stage of liver diseases, there are significant alterations of bile acids in plasma and urine (4). Thus, bile acids can be used as potential biomarkers for liver injury and dysfunction (5).

Rhein (4.5-dihydroxy anthraquinone-2-carboxylic acid, PubChem CID: 10168), a monoanthracene nuclear 1,8-dihydroxyanthraquinone derivative, is widely found in medicinal plants such as rhubarb, Sennae folium, Semen cassiae and Polygonum multiflorum which have laxative, choleretic and lipid-lowering functions (6). In the above plants, the amounts of rhein are approximately 4.7 mg/g (rhubarb) (7), 0.29 mg/g (Polygonum multiflorum) (8), 0.24 mg/g (Semen casssiae) (9), and 0.4 mg (Sennae folium) (10), suggesting that the amount of rhein in rhubarb is high. Recent studies have indicated that rhein has lipid-lowering anti-inflammatory, anti-tumor, and anti-hepatic fibrosis effects as wells as reduces blood glucose and improves renal interstitial fibrosis (11). These medical plants are broadly used all over the world. In China, approximately 10% (800) of the more than 8,000 proprietary Chinese medicines contain rhubarb (12). In the list of health care products published by the state food and drug administration of China in 2016, 66 kinds contained rhubarb, 253 kinds contained aloe, 440 species contained Semen casssiae, and 282 species contained Polygonum multiflorum. Most of them are used to help reduce blood lipids and lose weight (13). In addition to being used as a laxative in Europe, rhubarb is frequently used in food as a vegetable or in the production of desserts, jams and fruits (14).

Moreover, the cultivated area of rhubarb in some Nordic countries is approximately 60 hectares. Meanwhile, the rhubarb cultivation area in the United States and Canada is approximately 7 times the cultivated area of Europe (15). Rhubarb is widely popular in North America as an ingredient in pie (16). Rhubarb has the effects of promoting dampness and relieving jaundice (17), anti-inflammatory activity, kidney protection, preventing and treating high blood lipids and cholestasis, ameliorating fibrosis and hepatic encephalopathy, and promoting blood circulation and hemostasis (18). Anthraquinones are the active constituents of rhubarb, and they include rhein, emodin, chrysophanol, emodin methyl ether, and aloe-emodin (19); meanwhile, studies have indicated that rhein shows a higher bioavailability and is more easily absorbed and exposed than other anthraquinones (20, 21). However, the effects of repeated intake of rhein on liver function and bile acid metabolism are rarely reported. Therefore, the present study was designed to investigate the influences of different doses of rhein on liver and bile acid metabolism and provide information for the reasonable use of rhein.

## Results

### Physical effects of rhein

The body weights of rats in the 1,000 mg/kg group were notably reduced after treatment with rhein for 5 consecutive weeks compared with the control group (*p*<0.01, Fig. S1). In addition, the urine volume of rats in the 10 mg/kg and 30 mg/kg groups were remarkably increased compared with the control (data not shown). The relative liver weights were not different from the blank control. No diarrhea was found during the study period, and there were no other abnormal signs in the rats during the administration period.

### Multivariate regression analysis of bile acids in serum and livers

The LC/MS chromatograms of 14 serum and liver bile acids, including t-CBAs (Tauro-α-muricholic acid-T-α-MCA, taurocholic acid-TCA, taurohyodeoxycholic acid-THDCA, and taurodeoxycholic acid-TDCA), 5 glycine-conjugated bile acids (g-CBAs) (glycoursodeoxycholic acid-GUDCA, glycohyodeoxycholic acid-GHDCA, glycochenodeoxycholic acid-GCDCA, glycodeoxycholic acid-GDCA, and glycocholic acid-GCA), and 5 UCBAs (ursodeoxycholic acid-UDCA, chenodeoxycholic acid-CDCA, deoxycholic acid-DCA, β-muricholic acid-β-MCA, and cholic acid-CA) are shown in Fig. 1(a, c, e). The specific concentrations of serum and liver bile acids in the control and rhein treatment groups are shown in Table 1. Through a clustering analysis of rat serum and liver bile acid data, we can intuitively determine any differences between the rhein-treated or non-treated groups. The results of the principal component analysis showed that both serum and liver bile acid data were remarkably separated from the control group after rhein administration (Data not shown). An orthogonal partial least-squares discrimination analysis (OPLS-DA) revealed a good segregation between the treatment groups and the control (Fig. 2a, d). Although the 10 mg/kg and 30 mg/kg groups were not completely separated, they were still divided from the control. The 1,000 mg/kg rhein group was not only better distinguished from the control but also separated from the 10 mg/kg and 30 mg/kg groups(Fig. 2). These results indicate that the treatment of rats with rhein markedly altered the composition and levels of serum and liver bile acids. The analysis of the animal latent variable 1 (LV1) scores for both serum (Fig. 2b) and liver (Fig. 2e) showed that the bile acid levels in the rhein groups differed from the control. The variable importance in projection (VIP) values was used to assess potential markers (Fig. 2c, f), and a VIP value above 1.0 implied that the independent variable had a more important role in the interpretation of the dependent variable (22). Using this criterion, we screened TCA, THDCA, T-α-MCA, β-MCA, and GCA in serum and TCA, T-α-MCA, CA, DCA, and GCDCA in liver as potential markers.

**Table 1.** Concentrations of bile acids in serum and liver after rats were treated with rhein for 5 weeks. Data are presented as the means ± SD concentrations in serum measured using UPLC-MS/MS on 5 rats. \**p* < 0.05, \*\**p* < 0.01, \*\*\**p*<0.001, compared with the control group of the same bile acid.

|  | Control | Rhein |  |  |
| --- | --- | --- | --- | --- |
|  | - | 0.01 g/kg | 0.03 g/kg | 1g/kg |
| Serum-Male (ng/ml) |  |  |  |  |
| T-α-MCA | 383.6±105.53 | 241.60±111.27 | 146.40±31.94** | 826.80±256.84* |
| TCA | 18.62±3.59 | 25.92±12.84 | 13.16±7.94 | 38.86±16.74 |
| THDCA | 52.16±17.66 | 46.36±21.74 | 17.34±3.13* | 9.59±5.75** |
| TDCA | 121.44±68.99 | 96.58±47.50 | 113.88±71.61 | 6.01±3.51* |
| GUDCA | 2.66±0.80 | 2.22±1.13 | 1.32±1.55 | 0.56±0.34** |
| GHDCA | 14.74±4.59 | 25.68±13.82 | 14.13±10.80 | 39.90±20.29* |
| GCDCA | 27.5±11.91 | 30.14±21.74 | 7.15±2.95* | 11.02±3.37* |
| GDCA | 99.62±26.50 | 24.70±16.74** | 24.33±10.72** | 16.35±6.18** |
| GCA | 248.40±78.62 | 594.60±287.05 | 122.48±33.96* | 294.26±179.41 |
| UDCA | 387.4±133.98 | 107.94±54.82** | 95.08±18.45** | 213.46±112.03 |
| DCA | 723.2±179.79 | 91.60±36.62** | 127.00±22.55** | 56.08±34.09*** |
| β-MCA | 1,385.4±452.18 | 232.00±26.94** | 270.20±101.18** | 730.20±197.87* |
| CA | 3,132±1,231.91 | 281.80±90.75** | 963.80±365.11* | 3,740.00±1,789.06 |
| CDCA | 1,517.4±623.24 | 286.00±162.30** | 269.68±98.37* | 688.40±312.90* |
| Liver-Male ug/g |  |  |  |  |
| T-α-MCA | 8.11±2.97 | 24.94±6.33** | 29.49±10.89** | 60.88±23.79** |
| TCA | 48.95±15.76 | 36.33±15.75 | 38.80±13.97 | 110.87±32.40** |
| THDCA | 6.97±3.09 | 4.27±0.56 | 2.34±1.23* | 2.46±1.20* |
| TDCA | 25.86±5.17 | 11.67±3.24** | 9.93±5.07** | 0.41±0.25*** |
| GUDCA | 0.80±0.41 | 0.20±0.06* | 0.12±0.08* | 0.03±0.03* |
| GHDCA | 1.53±0.94 | 0.57±0.32 | 0.14±0.10* | 0.01±0.00* |
| GCDCA | 2.78±0.74 | 1.60±0.61* | 0.66±0.33** | 0.16±0.10** |
| GDCA | 11.22±2.41 | 2.29±1.74*** | 0.92±0.74*** | 0.02±0.01*** |
| GCA | 21.98±14.85 | 13.79±4.85 | 6.02±3.28 | 4.64±3.03 |
| UDCA | 0.23±0.16 | 0.11±0.04 | 0.15±0.03 | 0.02±0.01* |
| DCA | 1.11±0.63 | 0.23±0.30* | 0.63±0.41 | 0.16±0.25* |
| β-MCA | 2.07±0.69 | 0.44±0.15* | 0.57±0.26* | 0.14±0.08* |
| CA | 3.33±1.00 | 0.22±0.02** | 0.79±0.55** | 0.35±0.22** |
| CDCA | 0.26±0.23 | 0.05±0.01 | 0.10±0.04 | 0.07±0.02 |

**Fig. 1.**
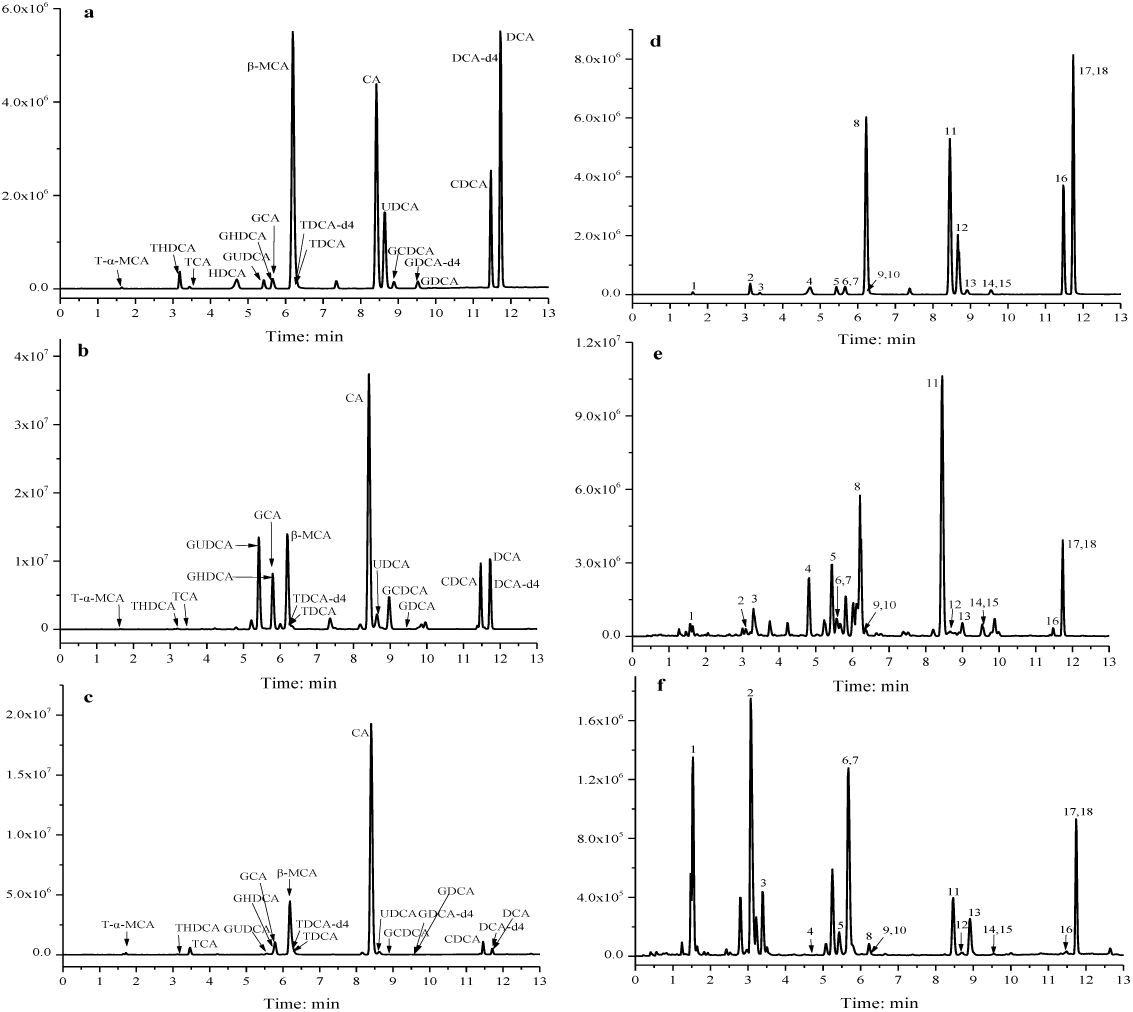
UPLC-MS/MS chromatogram of bile acids in serum and livers. Bile acid standards in serum samples (a), serum samples in the control group (b), serum samples in the 1,000 mg/kg group (c) and bile acid standards in the hepatic samples (d), hepatic samples in the control group (e), and hepatic samples in the 1,000 mg/kg group (f).

**Fig. 2.**
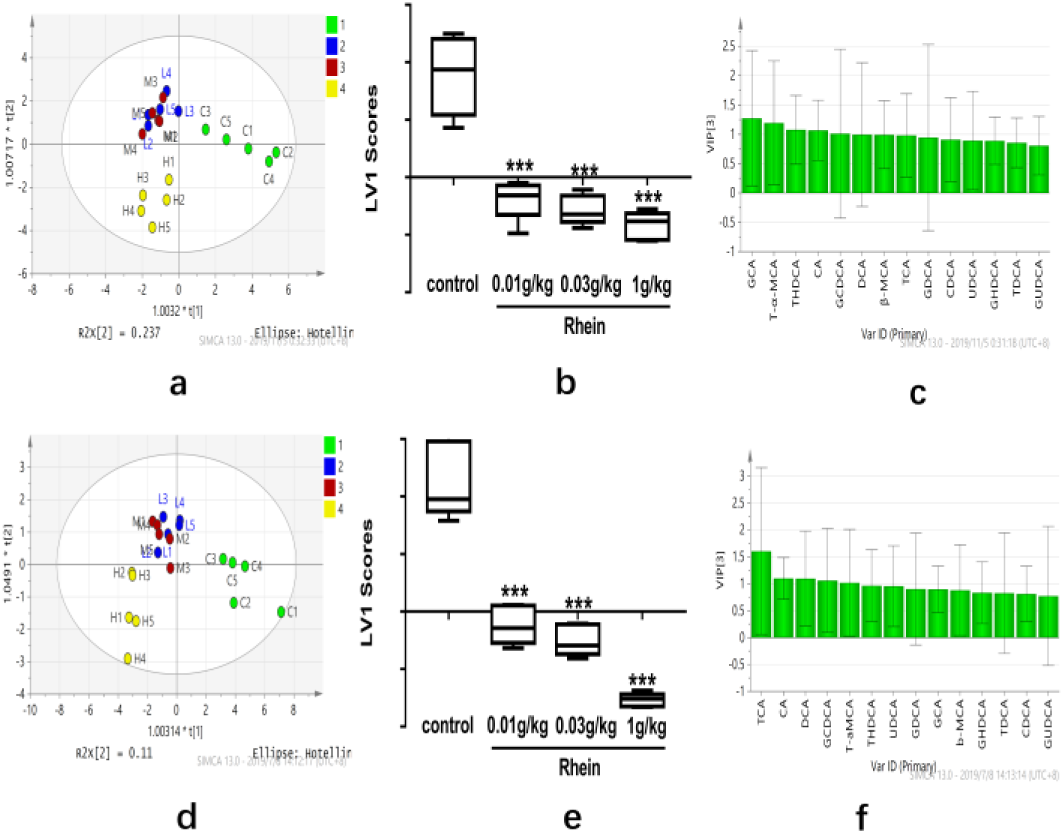
Multivariate data analysis of bile acid profiles in serum and liver. The OPLS-DA score plots demonstrated complete separation of the samples between groups in the serum (a) and liver (d). The green points represented the control, while the blue, red and yellow points represented the 10 mg/kg, 30 mg/kg and 1,000 mg/kg rhein group, respectively, as shown on the plots. The t1 scores in the serum (b) and liver (e) are shown, respectively, according to the OPLS-DA score plots. The VIP plots of OPLS-DA highlighted the discriminatory species in serum (c) and liver (f).

### Serum bile acids profiles after treatment with rhein

The amounts of 14 individual bile acids in serum were determined after the rats were treated with or without rhein (Table 1), and the contents of g-CBAs, t-CBAs, UCBAs and TBAs (sum of the 14 bile acids) were classified and analyzed (Fig. 3a, b). The results show that the level of UCBAs in serum was the highest. The composition ratio and contents of serum bile acids in rats were significantly affected by oral administration of rhein to rats for 5 weeks. In addition to reducing the level of GDCA, 10 mg/kg rhein had no significant effect on the overall contents of g-CBAs and t-CBAs. However, g-CBAs (especially GCDCA, GDCA and GCA, *p*<0.05 or *p* <0.01) as well as t-CBAs (especially T-α-MCA and THDCA, *p*<0.05 or *p*<0.01) were decreased after the rats were administered 30 mg/kg rhein. The dose of 1,000 mg/kg rhein can significantly reduce the amounts of 3 unconjugated bile acids (DCA, CDCA, b-MCA, *p*<0.05 or *p*<0.01) while slightly elevating the CA level compared with the control group. For this reason, 1,000 mg/kg rhein had no significant effect on the overall amounts of serum UCBAs. Meanwhile, the high dose of rhein affected the g-CBAs and t-CBAs contents remarkably. Two g-CBAs (GUDCA 375%↓, GCDCA 149.5%↓, and GDCA 509.2%↓) and two t-CBAs and GHDCA(170.7%↑) and two t-CBAs (T-α-MCA 115.5%↑, TCA 108.7%↑) were decreased and increased, respectively, compared with the control. The results demonstrate that different doses of rhein displayed a diverse influence on bile acid homeostasis, and 10 mg/kg and 30 mg/kg rhein may be beneficial to its therapeutic effect of reducing bile acids.

**Fig. 3.**
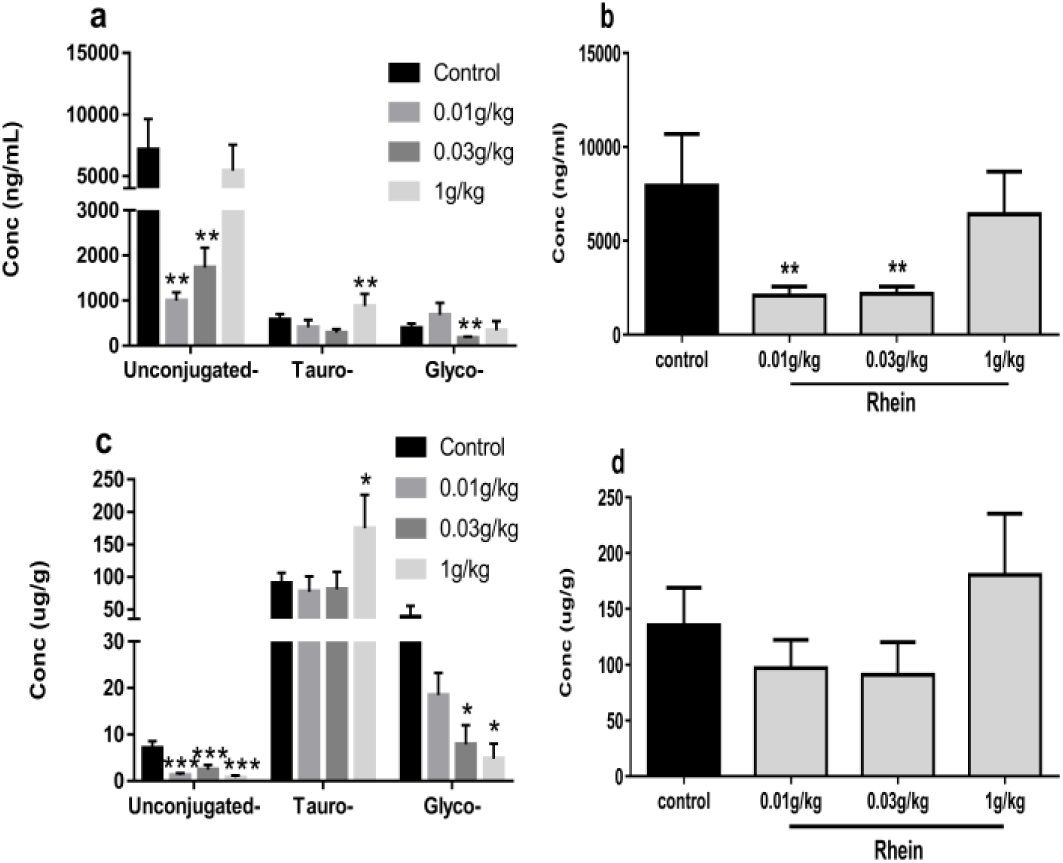
Alterations in the composition of bile acids in the serum and liver after the rats were treated with rhein. Serum concentrations of t-CBAs, g-CBAs and UCBAs (a) and concentrations of TBAs (b) in different groups. In addition, concentrations of t-CBAs, g-CBAs and UCBAs (c) and concentrations of TBAs (d) in the liver of different groups. Data are presented as M ± SD of 5 rats. \**p* < 0.05, \*\**p* < 0.01, compared with the control group.

### Liver bile acid profiles after treatment with rhein

The LC/MS chromatograms of 14 liver bile acids are shown in Fig. 1(b, d, f). From Fig. 3(c, d), we can find that the t-CBAs accounted for the majority of TBAs in all groups while the proportion of UCBAs was the lowest, which is contrary to that in the serum. The hepatic TBAs were slightly decreased after the rats were administered with 10 mg/kg and 30 mg/kg rhein; meanwhile, the TBA concentrations were elevated when the dose of rhein was increased to 1,000 mg/kg. Rhein notably decreased the contents of UDCA, DCA, β-MCA, CA, and CDCA in liver (Table 1). As a result, the overall levels of UCBAs in the liver were significantly decreased versus in the control (*p* < 0.05 or *p* < 0.01). The contents of 5 g-CBAs were reduced with oral administration of rhein to rats, showing a decreased trend with the elevation of the dose of rhein. For the t-CBAs, 10 mg/kg and 30 mg/kg rhein remarkably reduced the TDCA level and elevated the T-α-MCA level (*p* < 0. 01). The dose of 30 mg/kg rhein could also significantly reduced the level of THDCA versus the control. When the dose of rhein was up to 1,000 mg/kg, it not only had a more notable effect in decreasing the TDCA and THDCA levels and the increasing T-α-MCA level in the liver but also significantly elevated the content of TCA. These results suggest that large doses of rhein can increase TBAs levels in the liver, mainly t-CBAs.

### Impact of rhein on hepatic bile acid transport and gene expression

To investigate the mechanism of abnormal alterations of bile acids induced by rhein, we analyzed the gene expression of liver bile acid receptor farnesoid X receptor (FXR) and the transporters associated with bile acid synthesis, transport and excretion by quantitative real-time PCR. It can be seen from Fig. 4 that 10 mg/kg and 30 mg/kg rhein had no remarkable effect on the expression of BSEP, Na^+^-dependent taurocholic cotransporting polypeptide (NTCP) and multidrug resistance-associated protein 2 (Mrp2) mRNA in the liver, but it could upregulate the expression of FXR, Mrp3 and CYP7A1 mRNA after treating the rats with rhein for 5 weeks. When the dose of rhein was increased to 1,000 mg/kg, the expression of CYP7A1 mRNA could be continuously upregulated, but the expression of FXR mRNA was repressed, and there was a significant difference compared with the control group (*p* < 0.05). At the same time, 1,000 mg/kg rhein could suppress the expression of BSEP and NTCP mRNA, especially the expression of BSEP when compared with the control (*p* < 0.05). However, there was no significant effect on Mrp2 and Mrp3 mRNA expression. These results indicate that the effects of different doses of rhein lead to diverse effects on the gene expression involved in bile acid homeostasis.

**Fig. 4.**
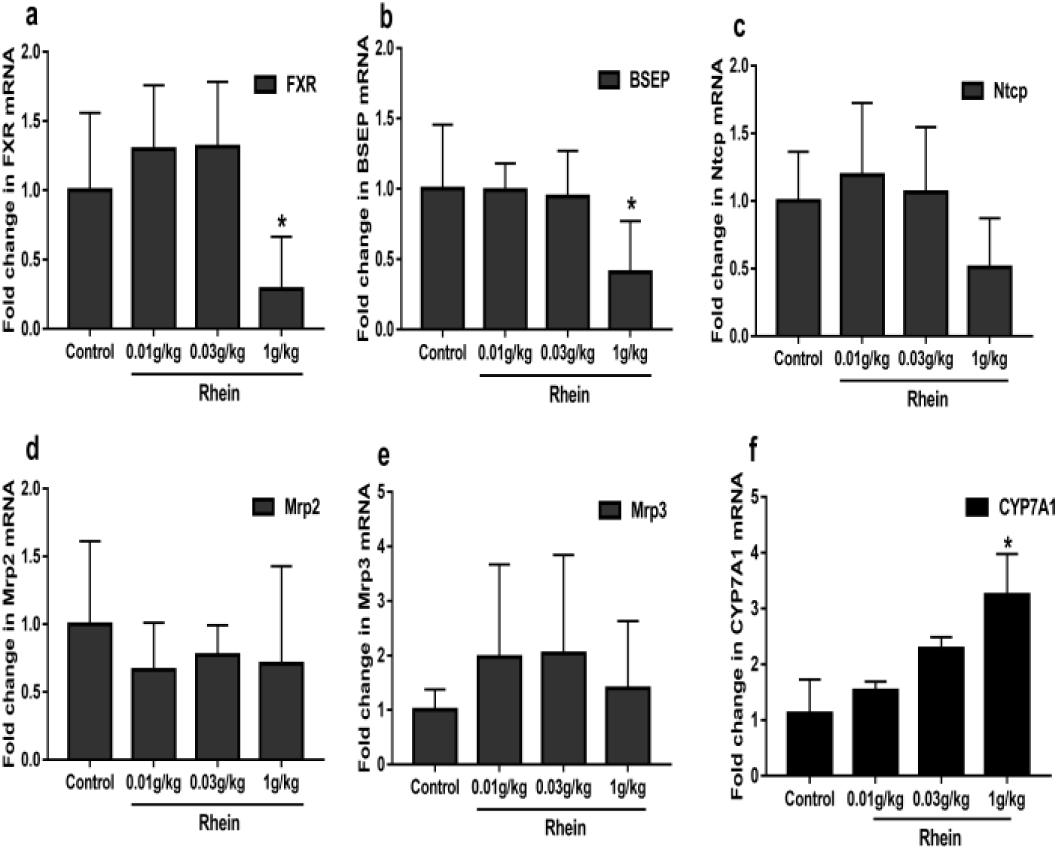
Expressions of genes involved in hepatic bile acid regulation. Quantitative real-time PCR analysis was performed to measure the expressions of genes in the livers, including FXR (a), BSEP (b), Ntcp (c), Mrp2 (d), Mrp3 (e) and CYP7A1 (f). Data are presented as M ± SD of 5 rats. \**p* < 0.05, compared with the control group.

### Serum biochemical parameters analysis

The serum biochemical indexes (Table 2) involved in liver function were investigated. The results show that there were no marked differences in serum ALT, AST and CHO levels after continuous intragastric administration of rhein for 5 weeks. The level of serum ALP in the 30 mg/kg group was lower than that in the control, but the change was not dose-dependent. The dose of 1,000 mg/kg rhein reduced the serum TBIL and γ-GT levels (*p* < 0. 05). However, the reduction of the above indicators did not have toxicological significance. The results imply that rhein has no significant toxicological effect on liver function and kidney function (data not shown).

**Table 2.**
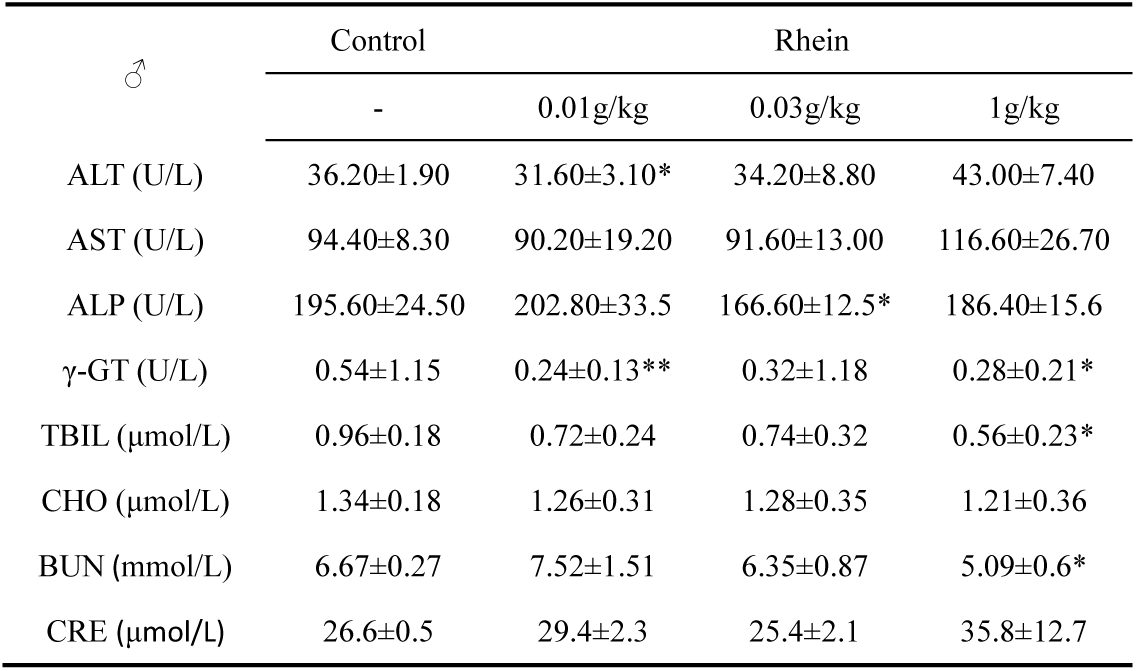
Serum biochemical values of ALT, AST, ALP, γ-GT, TBIL, CHO, BUN, CRE after rats treated with rhein for 5 weeks. Data are presented as means ± SD of 5 rats. \**p* < 0.05, compared with the control group.

### Histopathological examination of liver

Histopathological examination of the liver showed inflammatory infiltration and fat droplets at 1,000 mg/kg rhein. Furthermore, mild ductular proliferation was found in 1 of the 5 rats after it was given 1,000 mg/kg rhein for 5 continuous weeks (Fig. 5b). However, histological abnormalities were not observed in the control, 10 mg/kg and 30 mg/kg groups.

**Fig. 5.**
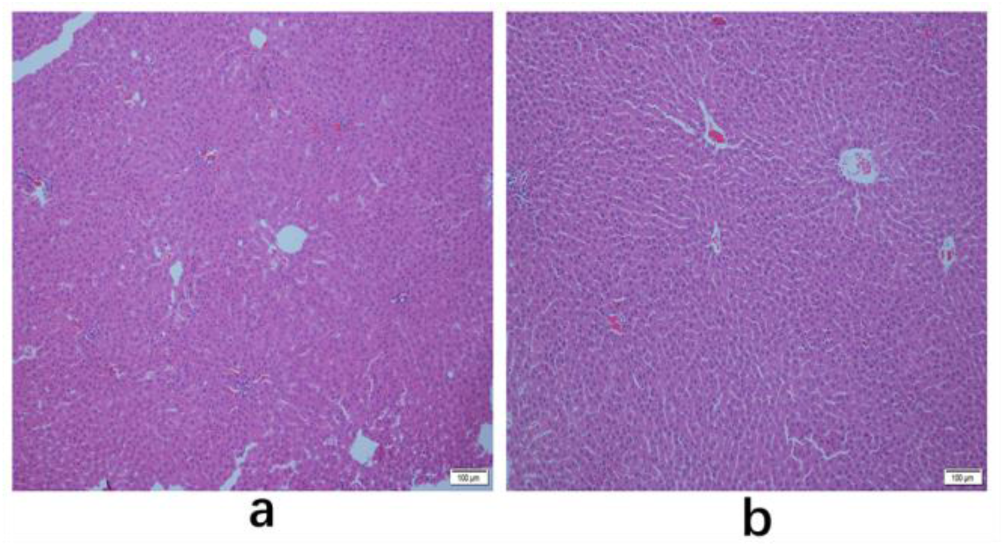
Histomorphological changes in the livers of rats with or without rhein treatment. Paraffin-embedded liver sections were stained with hematoxylin and eosin (HE). (a) Control group, (b) 1,000 mg/kg rhein group.

### Statistical analyses

The data are shown as the mean (M) ± standard deviation (SD). Statistical analysis was performed using one-way ANOVA with SPSS software (version 16.0). Bile acid data were also used to perform the OPLS-DA with SIMCA-P (version 12.0). Data were compared with the control group, and P values less than 0.05 were considered significant.

## Discussion

According to previous reports, rhein exerts significant anti-inflammatory effects (23), antioxidant effects (24), lipid-lowering effects (25), osteoarticular protection (26) and anti-diabetic activities (27) and improves renal interstitial fibrosis (28). The maximum daily dose of rhubarb for an adult is 15 g according to the 2015 edition of Chinese Pharmacopoeia (17); meanwhile, the content range of rhein in rhubarb is 0.47%-2.107% (7, 29). Therefore, the amount of rhein that an adult may ingest per day when taking 15 g rhubarb is 70.5 mg to 316 mg, which is equivalent to 7.4 mg/kg to 33 mg/kg in rats according to the dose conversion method between animals and humans (30). Considering the fluctuation of the amounts of rhein in medicinal materials, we chose 10 mg/kg rhein, close to the clinical equivalent dose, as the lowest dose in this study, and designated another 30 mg/kg and 1,000 mg/kg rhein (approximately equivalent to 3 times and 100 times of the clinical dose, respectively) to explore the possible influence of different doses of rhein on bile acid homeostasis.

In this study, treatment of rats with rhein for 5 weeks did not result in obvious injury to the liver, kidney and other important organs, and no toxicological effects were found in the blood biochemical indicators including AST, ALT, ALP, γ-GT, BUN and CRE, implying that rhein has low toxicity. The absolute concentrations of 14 endogenous bile acids in the serum and liver were obtained by UPLC-MS/MS with or without rhein administration; although it was observed that rhein notably altered the constitution and amounts of serum and liver bile acids, the analysis results of the serum and liver bile acids in each groups showed that rhein elevated the contents of t-CBAs and decreased the amounts of g-CBAs and UCBAs in the liver. Previous studies have reported that partial UCBAs such as CA, DCA, CDCA and β-MCA display cytotoxicity due to high hydrophobicity (31, 32). Taurine-conjugated bile acids (TCA, T-α-MCA) and glycine-conjugated bile acids, such as GCA, are less toxic (33). Therefore, the effect of rhein on bile acids is generally beneficial to the reduction of liver toxicity, and this advantage becomes evident when the level of rhein is close to clinical use (10 mg/kg and 30 mg/kg).

To elucidate the mechanism of the changes of bile acid induced by rhein, the gene expression associated with bile acid homeostasis was analyzed by quantitative real-time PCR. We know that bile acid synthesis takes place in the liver via two different pathways. The classical pathway is catalyzed by cholesterol 7 α-hydroxylase (CYP7A1), the rate-limiting enzyme (34) deciding the contents of bile acid synthesis and regulated by the FXR [35], to produce most bile acids. Meanwhile, FXR is an important regulator; it not only regulates bile acid synthesis but also plays an important role in bile acid transport and excretion (31, 36). Bile acids are secreted from the liver into the bile canaliculus via the canalicular bile salt export pump (BSEP) localized in the canalicular or apical domain of the hepatocyte plasma membrane (31), and a loss of BSEP leads to intrahepatic cholestasis (37). Multidrug resistance-associated protein 2 (Mrp2), located in the canalicular membrane, can also induce bile acids to be secreted from the liver into bile (2).

Sodium taurocholate cotransporting polypeptide (NTCP), localized on the outside of the basolateral membrane, largely accounts for the uptake of conjugated bile acids (> 80%) (38). Meanwhile, multidrug resistance-associated protein 3 (Mrp3), localized at the basolateral membrane of hepatocytes, excretes bile acids into the blood (2). Our study showed that rhein influences the synthesis, metabolism and excretion of bile acids by affecting the expression of bile acid transporter genes. The multivariate statistical analysis demonstrated a notable distinction between the rhein group and the control, and the main reason for this remarkable distinction was the variation in TCA and T-α-MCA levels in the serum and liver. The dose of 1,000 mg/kg rhein notably repressed the expression of hepatic messenger RNA of FXR while remarkably elevating the expression of CYP7A1 mRNA, leading to increased bile acid synthesis. On the other hand, 1,000 mg/kg rhein also downregulated the expression of BSEP mRNA, resulting in decreased secretion of bile acid to the bile duct and thus causing an increase in the total amount of bile acid in the liver (Fig. 3d). However, excessive amounts of bile acids in hepatocytes may cause damage to hepatocytes. It should be noted that no significant liver damage was observed at the dose of 1,000 mg/kg rhein during our study period, and this may be related to the increase of t-CBAs (TCA, T-α-MCA) that are less toxic. It is important to note that high doses of rhein have the potential to increase risk; thus, taking excessive doses of rhein over a long period should be avoided. However, 1,000 mg/kg rhein is much higher than the clinical dose (approximately 100 times the clinical dose), and the chance of ingesting such a high dose of rhein is very low. The 10 mg/kg and 30 mg/kg rhein doses used in this study are close to or slightly higher than the clinical dose (clinical dose and 3 times of clinical dose, respectively), potentially resulting in slight upregulation of the expression of liver FXR mRNA and MRP3 mRNA; this in turn inhibits the synthesis of excessive amounts of hepatic bile acids and induces bile acids to be secreted, leading to reduced liver bile acids, especially the UCBAs (Fig. 3c) that are relatively high toxic. This is beneficial to the hepatocytes. No significant abnormalities or hepatoxicity were found in either the 10 or 30 mg/kg groups. Therefore, rhein is safe to use at a reasonable dosage and timing.

## Materials and methods

### Chemicals and reagents

Rhein (Fig.6) was purchased from Beijing Saibaicao Technology Co., Ltd., and its purity was more than 98% according to high-performance liquid chromatography. The chemical structure of rhein is shown in Figure S1. All reference bile acids purchased were of high purity. TCA (PubChem CID: 6675) was purchased from Cayman Chemical Company (Ann Arbor, Michigan, US). T-α-MCA (PubChem CID: 101657566), TDCA (PubChem CID: 2733768), GUDCA (PubChem CID: 12310288), GHDCA (PubChem CID: 114611) and β-MCA (PubChem CID: 5283853) were purchased from Toronto Research Chemicals (Toronto, Canada). GCDCA (PubChem CID: 12544), GDCA (PubChem CID: 3035026) and GCA (PubChem CID: 10140) were purchased from Nanjing Shenglide Technology Co., Ltd. (Nanjing, China). THDCA (PubChem CID: 119046), CA (PubChem CID: 221493), UDCA (PubChem CID: 31401), HDCA (PubChem CID: 5283820), CDCA (PubChem CID: 10133) and DCA (PubChem CID: 222528) were purchased from the National Institutes for Food and Drug Control (NIFDC, Beijing, China). Deuterated internal standards DCA-d4, GDCA-d4, TDCA-d4 were obtained from Cambridge Isotope Laboratories (Tewksbury, US).

**Fig. 6.**
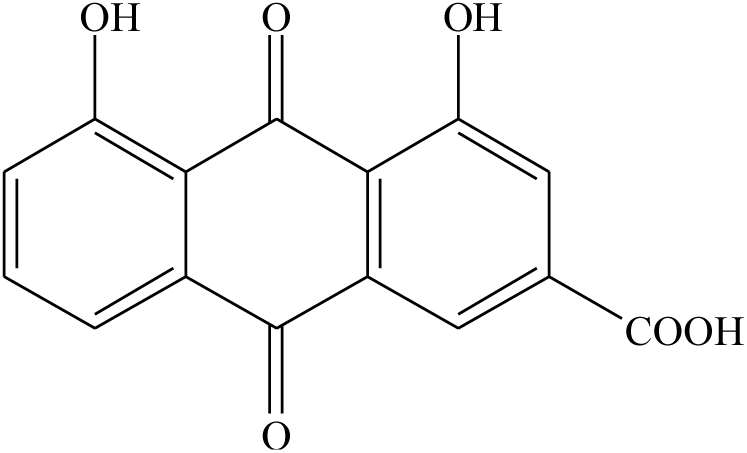
Chemical structure of rhein.

### Animals and experimental procedure

Twenty male Sprague-Dawley rats at 6-8 weeks of age with body weights ranging from 70–90 g were obtained from Vital River Laboratory Animal Technology Co., Ltd. (Beijing, China). They were maintained in a controlled environment (temperature 23 ± 3 °C, humidity of 40∼70%, 12 h light/dark cycle). The rats were allowed free access to standard food and pure water. The animals were randomly divided into 0, 10, 30, and 1,000 mg/kg rhein groups. The rats were dosed by a gastric gavage once a day for 5 consecutive weeks while the rats in the control group were administered an equal volume of water. The rats were anesthetized with sodium phenobarbital by intraperitoneal injection. Blood samples were collected from the abdominal aorta by syringe under the condition of fasting 16 h, and then serums and livers were obtained. The serum samples were used for the analysis of bile acids and biochemical indexes. The livers were used for pathological examination, the analysis of gene expression and the detection of bile acids. The experiment was performed according to the ethical guidelines for the use of laboratory animals. This study was approved by the Committee on the Ethics of Animal Experiments of the Institute of Chinese Materia Medica, China Academy of Chinese Medical Sciences (Permit Number: 20152018).

### Analysis of bile acids in serum and liver by LC/MS Conditions of chromatography and mass spectrometry

The mobile phase comprised 0.01% formic acid in water (A) and 0.01% formic acid in acetonitrile (B). The gradient of A was as follows: 75–60% A (0– 9 min), 60–35% A (9–13 min), 35–1% A (13–16 min), and 1-75% A (16–18 min). The mass spectrometer was operated in negative mode with the multiple reaction monitoring (MRM) function for the quantification (39), and more details on the MRM conditions are shown in Supplementary Table S1. The total chromatographic operation was divided into several periods. The ion dwell time and transition for the bile acids were set reasonably. The mass spectrometer was operated with the source and desolvation temperatures set at 120 °C and 350 °C, respectively. The curtain gas was 40 psi; the ion spray voltage was −4500 V; the probe temperature was 600 °C; the collision gas was medium; and ion source gas 1 and ion source gas 2 were all 40 psi.

### Sample preparation for LC/MS analysis

For the serum samples, Oasis-HLB solid-phase extraction (SPE) columns were used for sample extraction. Deuterated standards of CA, GCA, and TCA were used as internal standards (ISs). First, 100 µL serum samples were spiked with 10 µL of the working solution of ISs (1 µg/mL) and 890 µL 0.01% formic acid water, vortexed, then loaded onto SPE columns preactivated with 1 mL MeOH, followed by 2 mL 0.01% formic acid water. Then, the loaded cartridges were washed with 1 mL 10% MeOH, 1 mL 0.05% formic acid, and 1 mL 45% methanol and eluted with 3 mL MeOH. The eluate was evaporated to dryness by N2 at 37 °C and reconstituted in 100 µL MeOH, and then 1 µL was injected into the AB SCIEX QTRAP 6500 LC/MS system. For each 100 mg of liver, 600 µL of cold methanol and 10 µL of the 1 µg/mL working solution of ISs were added.

Then, the liver tissues were homogenized. Tubes were centrifuged at 6,000 rpm/min for 10 min at 4 °C, and the supernatants were transferred to clean tubes. A second BA extraction was performed using 400 µL of cold methanol. Finally, the two extraction supernatants were combined and centrifuged at 15,600 rpm/min for 10 min at 4 °C. Then, the supernatants were evaporated to dryness by N2 at 37 °C and reconstituted in 100 µL MeOH, and 1 µL was injected into the AB SCIEX QTRAP 6500 LC/MS system.

### Calibration and quality control standard preparation

The bile acid standards were dissolved in methanol to obtain 1 mg/mL of BA stock solution or IS stock solution. The individual BA stock solutions were combined and diluted to obtain the mixed working standard solution (50 µg/mL) or working ISs solution (1 µg/mL) of bile acids. Eight calibration standard solutions ranging from 2 to 10,000 ng/mL were obtained by serially diluting the working standard solution into charcoal-stripped serum. Quality control (QC) samples were acquired in the same manner at 100, 1,000, and 10,000 ng/mL in charcoal-processed serums. The calibration standards and QC standards were then disposed of according to the sample preparation process depicted above. The linear regression parameters obtained for serum bile acids and liver bile acids are shown in Supplementary Table S2 and Table S3, respectively. Five replicates of each QC point were analyzed to evaluate the intra- and interday accuracy and precision (Supplementary Table S4 and Table S5). This process was repeated 3 times over 3 days to evaluate the interday accuracy and precision. The extraction recoveries of 14 bile acids are shown in Supplementary Table S3.

### Individual bile acid analysis

The individual bile acid (BA) analysis was performed by LC/MS. The mass spectrometer was an AB SCIEX 6500 QTrap equipped with an Acquity UPLC system operating in negative mode. Chromatographic separation of the bile acid was performed on an ACQUITY UPLC BEH column (2.1 × 100 mm, 1.7 μm) (Waters Corp., Milford, US) maintained at 35°C at a flow rate of 500 µL/min. The deuterated standards d4-TCA, d4-GCA, and d4-CA were used as ISs for TCA, GCA, and CA, respectively. Peak integration and quantification were performed using Analyst 1.6.2 software. Each standard curve for bile acids was built by plotting the ratio of the BA peak area to its deuterated standard peak area versus the concentration. The slope and y-intercept were computed using a linear curve fit with 1/x^2^ weighting. The concentrations of individual BA in the samples were computed according to the regression line. Data were processed with SIMCA-P software, version 12.0.

### Biochemical assay and histopathology

Aspartate aminotransferase (AST), alanine aminotransferase (ALT), alkaline phosphatase (ALP), γ-glutamyltransferase (γ-GT), cholesterol (CHO), total bilirubin (TBIL), blood urea nitrogen (BUN) and creatinine (CRE) were examined using a TBA-40FR automatic biochemistry analyzer (Toshiba, Japan). Histopathological examination of the liver was performed in all rats. Organ samples were fixed with neutral buffered formalin, embedded in paraffin, sectioned, and stained with HE. The histomorphology was examined under a light microscope (Olympus, Japan).

### Quantitative Real-time PCR analysis

Total hepatic RNA was prepared by a total RNA kit (OMEGA, Georgia, USA) in accordance with instructions. An aliquot of 1 µg RNA was applied for reverse transcription with oligo-dT primer (TOYOBO, OSAKA, Japan). Quantitative real-time PCR was performed using the Roche 480 instrument (Roche, Mannheim, Germany) and SYBR Green PCR Master Mix (Roche, Mannheim, Germany) for the subsequent genes with the corresponding primers (Sangon Biotech, Beijing, China) (Supplementary Table S6). Quantification was conducted using the ΔΔCT method. The quantity of messenger RNA was normalized with the internal standard GAPDH.

### Statistical analysis

The results are shown as the mean ± SD. Data were analyzed using SPSS 16.0 software, and differences between groups were analyzed by Student’s *t* test. Bile acid data were also determined by OPLS-DA using SIMCA-P 12.0 software. The data of the rhein treatment groups were compared with the control. Values significantly different from the control are indicated as \*\**p* <0.01 and \**p* <0.05.

## Conclusion

In conclusion, 1,000 mg/kg rhein notably suppressed the expression of FXR mRNA, while remarkably elevating the expression of CYP7A1 mRNA, leading to increased hepatic bile acid synthesis. On the other hand, 1,000 mg/kg rhein also downregulated the expression of BSEP mRNA, resulting in decreased secretion of bile acid to the bile canaliculus and thus causing an increase in the total amounts of bile acid in the liver. Ten and 30 mg/kg rhein can reduce the synthesis and promote bile acid clearance by upregulating the expression of FXR mRNA and Mrp3 mRNA, potentially resulting in the decrease in bile acid, which may be the mechanism of rhein in clinical application. Consequently, we believe that rhein is safe to use at a reasonable dosage and timing. However, However, the high dose of rhein increased the TBAs levels, which may result in the accumulation of bile acids in the liver; thus, the T-α-MCA and TCA levels should be monitored when taking rhein in high doses over a long period.

